# Activated Oncostatin M signaling drives cancer-associated skeletal muscle wasting

**DOI:** 10.1101/2023.01.26.525658

**Authors:** Aylin Domaniku, Samet Agca, Batu Toledo, Sevval Nur Bilgic, Aynur Erkin Kashgari, Serkan Kir

**Affiliations:** Department of Molecular Biology and Genetics, Koc University, Istanbul 34450, Turkey

**Keywords:** Cancer cachexia, skeletal muscle atrophy, muscular dystrophy, Oncostatin M, JAK/STAT3 signaling

## Abstract

Progressive weakness and muscle loss are associated with multiple chronic conditions including muscular dystrophy and cancer. Cancer-associated cachexia, characterized by dramatic weight loss and fatigue, leads to reduced quality of life and poor survival. Inflammatory cytokines have been implicated in muscle atrophy, however, available anti-cytokine therapies failed to prevent muscle wasting in cancer patients. We previously reported that muscle-specific deletion of the Oncostatin M (OSM) receptor (OSMR) preserved muscle mass and function in tumor-bearing mice. Here, we show that OSM is a potent inducer of muscle atrophy. OSM triggers cellular atrophy in primary myotubes utilizing the JAK/STAT3 pathway. Identification of OSM targets by RNA sequencing revealed the induction of various muscle atrophy-related genes, including *Atrogin1*. OSM overexpression in mice caused muscle wasting while the neutralization of circulating OSM protected from tumor-driven loss of muscle mass and function. Our results indicate that activated OSM/OSMR signaling drives muscle atrophy, and the therapeutic targeting of this pathway may be useful in preventing muscle wasting.

## Introduction

Skeletal muscle atrophy is characterized by the loss of muscle mass due to excessive protein catabolism and the consequent poor muscle strength (Cohen et al., 2015). Muscle loss is associated with multiple conditions, including aging (i.e., sarcopenia), organ failures, immune system complications, and muscular dystrophies (Cohen et al., 2015; Mercuri et al., 2019). Muscle wasting is also a hallmark of cachexia syndrome linked to chronic diseases such as cancer, kidney disease, and heart failure (Argiles et al., 2018). Cachexia-led progressive atrophy of muscle and adipose tissues causes dramatic weight loss, which is associated with poor quality of life due to frailty and restrained daily activity. Cachexia is particularly common among cancer patients and is the direct cause of approximately 20% of all cancer deaths (Argiles et al., 2014). It leads to a poor response to chemotherapy, often prevents patients from receiving further therapies, and negatively influences survival. With no effective treatment to block muscle wasting, cachexia has remained a significant unmet medical need (Dolly et al., 2020).

Systemic inflammation has long been linked to cancer cachexia, and pro-inflammatory cytokines TNFα, IL-1, and IL-6 have been suggested as causal agents (Webster et al., 2020). While TNFα has many systemic effects, its role in muscle wasting remains unclear, and the lack of a direct impact on protein breakdown in isolated muscles was reported (Kettelhut et al., 1987; Kettelhut and Goldberg, 1988). Anti-TNFα therapies failed to prevent muscle atrophy in patients with advanced cancer cachexia (Jatoi et al., 2007; Jatoi et al., 2010). IL-1 and IL-6 are upregulated in animal models of cancer cachexia, and targeting IL-6 ameliorates muscle loss in cachectic mice (Cohen et al., 2015; Strassmann et al., 1992). Clinical trials using anti-IL-1 and anti-IL-6 therapies showed promising results, however, a satisfactory effect on skeletal muscle mass was not attained (Bayliss et al., 2011; Marceca et al., 2020). To design novel targeted therapies, a better understanding of the link between inflammatory mediators and muscle atrophy is needed.

Investigating muscle wasting in a murine model of cancer cachexia, we identified Oncostatin M (OSM) as a potential mediator of inflammatory responses in skeletal muscle. OSM is a member of the IL-6 family of cytokines and has crucial functions in cell growth, differentiation, and inflammation (Hermanns, 2015). OSM was originally identified for its ability to inhibit tumorigenesis (Zarling et al., 1986). However, it modulates a variety of biological processes that are cell type-dependent, including liver development, hematopoiesis, and bone metabolism (Richards, 2013). Elevated OSM levels have been observed in cancer and inflammatory diseases in humans (Hermanns, 2015; Richards, 2013). OSM signals through its receptor, OSMR, and the receptor subunit GP130. *Osmr* gene itself is a transcriptional target of OSM (Blanchard et al., 2001; Stephens et al., 2018). Upon detecting elevated *Osmr* mRNA levels in muscles of tumor-bearing cachectic mice, we investigated the role of OSM/OSMR signaling in wasting. We found that OSM is a potent inducer of muscle atrophy in cultured primary myotubes and in muscle tissue of mice. In fact, muscles-pecific OSMR-knockout mice are resistant to tumor-driven muscle wasting (Bilgic et al., 2023). We showed that neutralization of OSM by a specific antibody preserves muscle mass and function in tumor-bearing mice. Our findings argue that OSM/OSMR signaling is a crucial driver of muscle loss.

## Results

### OSM promotes cellular atrophy in cultured primary myotubes

Utilizing the murine Lewis Lung Carcinoma (LLC) model of cancer cachexia, we detected highly elevated mRNA levels of the *Osmr* gene in atrophying muscles (Figure 1A). Similarly, the expression of E3 ubiquitin ligase genes *Atrogin1* and *MuRF1* whose protein products are well recognized to activate protein breakdown associated with muscle atrophy was also elevated. To test whether enhanced OSMR activity contributes to the muscle atrophy process, we isolated mouse primary myoblast cells and differentiated them into myotubes. Treatment of the primary myotubes with a recombinant OSM protein induced expression of *Atrogin1* without altering *MuRF1* levels (Figure 1B). OSM treatment also increased mRNA levels of its receptor *Osmr* along with mRNA levels of the downstream components of cytokine signaling; Janus Kinase 2 (*Jak2*) and Suppressor of Cytokine Signaling 3 (*Socs3*) (Figure 1B). Furthermore, OSM-treated myotubes exhibited reduced diameter, a sign of cellular atrophy (Fig 1C). OSM promoted a stronger atrophy-inducing effect as myotube diameters were reduced more significantly compared to other IL-6 family cytokines implicated in muscle atrophy; IL-6 and LIF (Figures 1C and 1D). To compare the atrophic effects of these cytokines, we tested protein levels of myosin heavy chain (MyHC) in myotubes. OSM administration reduced MyHC in primary myotubes indicating enhanced protein breakdown (Figure S1A). Immunofluorescently-labeled MyHC signal was also significantly reduced in OSM-treated myotubes where a more pronounced atrophy phenotype was observed compared to cells treated with IL-6 and LIF (Figures S1B-S1C).

**Figure 1.**
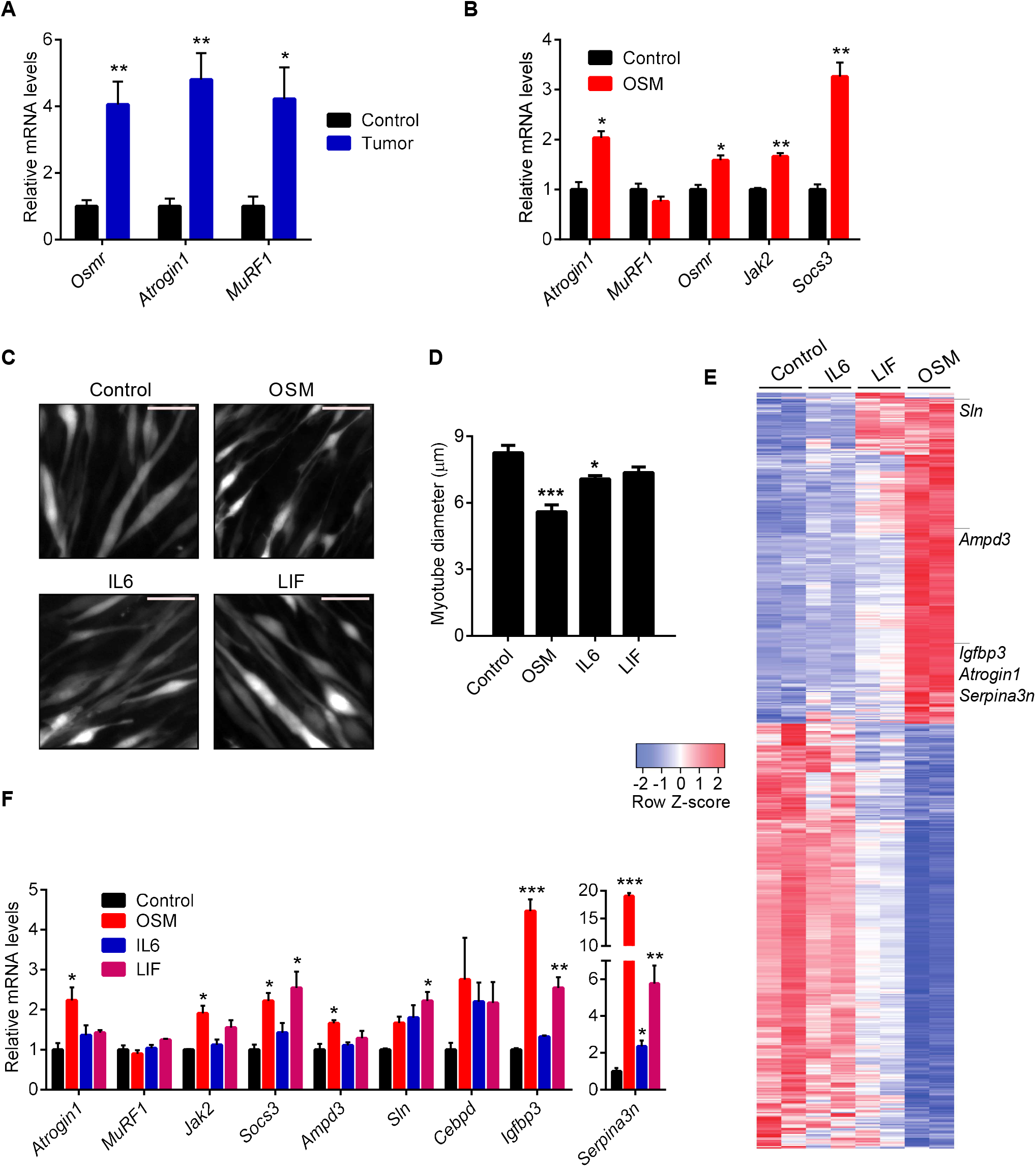
OSM promotes cellular atrophy in cultured primary myotubes. (A) C57BL/6 mice inoculated with LLC cells were sacrificed 16 days later and changes in gene expression of gastrocnemius muscle were determined by RT-qPCR (n = 5 for the control group and n = 7 for the tumor group). (B) Mouse primary myotubes were treated with recombinant OSM (250 ng/ml) for 48 hr. Changes in gene expression were determined by RT-qPCR (n = 3 for each group). (C and D) Mouse primary myotubes were transduced with a GFP adenovirus. Cells were treated with recombinant OSM, IL6 or LIF (each 250 ng/ml) for 48 hr and then visualized under the fluorescence microscope. Scale bar is 50 μm (C). Average myotube diameter was measured (n = 4 for each group) (D). (E and F) Mouse primary myotubes were treated with recombinant OSM, IL6 or LIF (each 250 ng/ml) for 48 hr. Gene expression profiles were analyzed by RNA sequencing. Heatmap of significant genes up- or down-regulated more than 2-fold is shown (n = 2 for each group) (E). Changes in gene expression were determined by RT-qPCR (n = 3 for each group) (F). Data are presented as mean ± SEM. Statistical analysis was conducted using two-tailed t-test (A, B) and one-way ANOVA with Tukey’s post-hoc test (D, F). \**p* < 0.05, \*\**p* < 0.01, \*\*\**p* < 0.001, compared with the control group.

We investigated the impact of these cytokines on the global gene expression profiles of mouse primary myotubes using RNA sequencing. Expression profiles of OSM-treated myotubes varied the most from the control group as evidenced by principal component analysis (Figure S2A). The highest number of genes with significant differential expression was detected in the OSM treatment group (Figures S2B-S2C). OSM appeared to have a larger footprint on the transcriptome of myotubes as visualized in the heatmap of the differentially expressed genes (Figure 1E). A list of the genes with significant expression changes is presented in Table S1 and the datasets can be found in Gene Expression Omnibus (GEO) database with accession number GSE222208. Pathway enrichment analysis demonstrated that genes upregulated by OSM include targets of JAK/STAT signaling involved in hallmark pathways such as inflammatory response, cytokine signaling and hypoxia whereas genes down-regulated by OSM are mostly related to cell cycle regulation (Figure S1D). From the analysis of RNA sequencing data, several muscle atrophy-related genes, including *Atrogin1 (Fbxo32), Ampd3, Sln, Cebpd, Igfbp3*, and *Serpina3n*, were found to be upregulated (Figure 1E and Table S1). We tested mRNA levels of these genes in mouse primary myotubes treated with OSM, IL-6 or LIF. In line with the impact on the myotube appearance, OSM treatment stimulated a significantly larger increase in *Atrogin1* levels (Figure 1F). Similarly, more pronounced changes in the expression levels of *Ampd3, Igfbp3* and *Serpina3n* were detected in response to OSM treatment (Figure 1F). Our findings argue that OSM is a potent inducer of cellular atrophy in primary myotubes and it significantly alters myotube gene expression.

### OSM utilizes JAK/STAT3 signaling to elicit its effects in myotubes

OSM/OSMR signaling is known to activate STAT transcription factors and the involvement of the JAK/STAT pathway in the atrophying muscle tissue was previously reported (Bonetto et al., 2012; Richards, 2013). Therefore, we investigated the effect of OSM administration on the phosphorylation and activation of the JAK/STAT signaling components. Treatment of mouse primary myotubes with recombinant OSM induced the phosphorylation of JAK2, STAT1, STAT3, and STAT5 (Figure 2A). Similar responses were obtained when myotubes were treated with recombinant LIF while IL6 treatment elicited milder effects. OSM-induced phosphorylation events were completely blocked when myotubes were also treated with JAK1/2 kinase inhibitor Ruxolitinib (Figure 2B). In accordance, OSM-led changes in mRNA levels of atrophy-related genes were also reversed by Ruxolitinib treatment, arguing an indispensable role for JAK kinases at downstream of the OSM signaling (Figure 2C). Activation of the NFκB signaling was also previously reported in the atrophying muscles (Cai et al., 2004). However, treatment of primary myotubes with OSM did not alter the phosphorylation of NFκB signaling elements, including IκB, p105-NFκB and p65-RelA (Figure S3A). In addition, the processing of p105-NFκB and p100-NFκB2 into p50-NFκB and p52-NFκB2, respectively, was unaffected (Figure S3B).

**Figure 2.**
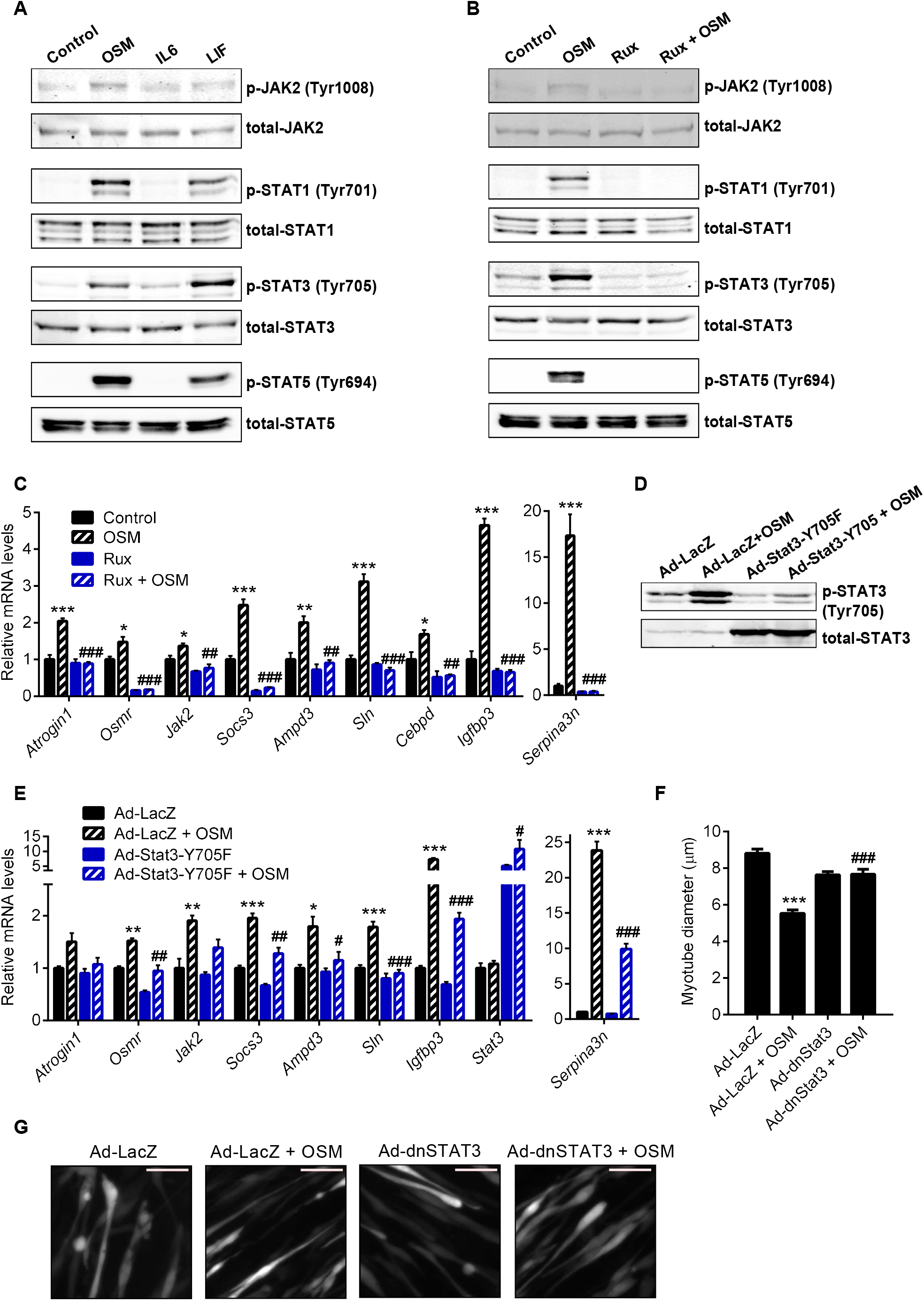
OSM utilizes JAK/STAT3 signaling to elicit its effects in myotubes. (A) Mouse primary myotubes were treated with recombinant OSM, IL6 or LIF (each 250 ng/ml) for 10 min. Protein levels were determined by western blotting. (B) Mouse primary myotubes were treated with Ruxolitinib (Rux; 2 μM) for 30 min and then recombinant OSM (250 ng/ml) was added for 10 min. Protein levels were determined by western blotting. (C) Mouse primary myotubes were treated with Ruxolitinib (Rux; 2 μM) and recombinant OSM (250 ng/ml) for 48 hr. mRNA levels were determined by RT-qPCR (n = 3 for each group). \**p* < 0.05, \*\**p* < 0.01, \*\*\**p* < 0.001 compares differences between Control and OSM groups. ##*p* < 0.01, ###*p* < 0.001 compares differences between OSM and Rux + OSM groups. (D-G) Mouse primary myotubes were transduced with LacZ or dominant-negative (dn) Stat3-Y705F expressing adenoviruses and treated with recombinant OSM (250 ng/ml) for 48 hr. Protein levels were determined by western blotting (D). mRNA levels were tested by RT-qPCR (n = 3 for each group) (E). Cells were also transduced with a GFP adenovirus for fluorescence imaging. Average myotube diameter was measured (n = 4 for each group) (F). Myotubes were visualized under the fluorescence microscope. Scale bar is 50 μm (G). \**p* < 0.05, \*\**p* < 0.01, \*\*\**p* < 0.001 compares differences between Ad-LacZ and Ad-LacZ + OSM groups. #*p* < 0.05, ##*p* < 0.01, ###*p* < 0.001 compares differences between Ad-LacZ + OSM and Ad-Stat3-Y705F + OSM groups. Data are presented as mean ± SEM. Statistical analysis was conducted using one-way ANOVA with Tukey’s post-hoc test.

We next investigated the activation of STAT transcription factors in muscle tissue of LLC-tumor bearing mice. In line with previous reports, tumors induced STAT3 phosphorylation in muscle (Bonetto et al., 2012; Bonetto et al., 2011; Silva et al., 2015) while the expression of *Atrogin1, MuRF1, Ampd3, Serpina3n* and *Cebpd* also increased (Figures S3C-S3E). Muscle-specific STAT3 depletion was previously shown to attenuate tumor-driven muscle wasting and STAT3 involvement in *Atrogin1* transcription was reported (Bonetto et al., 2012; Silva et al., 2015). Therefore, we tested the role of STAT3 in OSM-induced atrophy using a dominant-negative STAT3 isoform. STAT3-Y705F mutant was overexpressed in mouse primary myotubes by adenoviral delivery. The dominant-negative form blocked OSM-driven phosphorylation and activation of endogenous STAT3 protein (Figure 2D). Importantly, OSM-induced changes in the expression of atrophy-related genes were suppressed by the overexpression of dominant-negative STAT3 (Figure 2E). These myotubes were also resistant to OSM-driven cellular atrophy as myotube diameter did not change (Figures 2F and 2G). These findings indicate that OSM utilizes the JAK/STAT3 signaling to promote the expression of atrophy genes and the resultant cellular atrophy in primary myotubes.

### OSM overexpression causes muscle atrophy in mice

We investigated the potential of OSM to promote muscle atrophy *in vivo* by overexpressing it in the tibialis anterior (TA) muscles of mice. For this purpose, we generated an adenoviral vector expressing mouse OSM. When mouse primary myotubes were transduced with the OSM adenovirus, they were induced to undergo atrophy as evidenced by a decrease in their diameter (Figures S4A and S4B). Overexpression of OSM in primary myotubes also stimulated the expression of *Osmr* and muscle atrophy-related genes, including *Atrogin1* (Figure S3C). Next, we administrated the OSM adenovirus into the TA muscle unilaterally while the contralateral TA muscle was transduced with a control LacZ adenovirus. The weight of TA muscles fell significantly 7 days after the transduction with Adeno-OSM while gastrocnemius muscles from the same animals were unaffected (Figure 3A). Hematoxylin and eosin (H&E) staining of TA tissue sections demonstrated a significant drop in muscle fiber cross-sectional area in response to Adeno-OSM (Figures 3B and 3C). An increase in the frequency of fibers with a smaller cross-sectional area was detected (Figure 3D). The overexpression of OSM in muscle tissue induced mRNA levels of *Osmr*, and the atrophy-related genes, *Atrogin1, Ampd3, Cebpd, Igfbp3, Sln* and *Serpina3n* (Figure 3E). OSM overexpression also increased total protein levels of Atrogin1, MURF1, STAT1, STAT3 and STAT5 in muscle tissue (Figures 3F and 3G). However, phosphorylation of only STAT3 was stimulated by OSM relative to the total protein levels (Figures 3F and 3G). These results argue that activation of the OSM/OSMR signaling is able to promote atrophy in muscle tissue *in vivo*.

**Figure 3.**
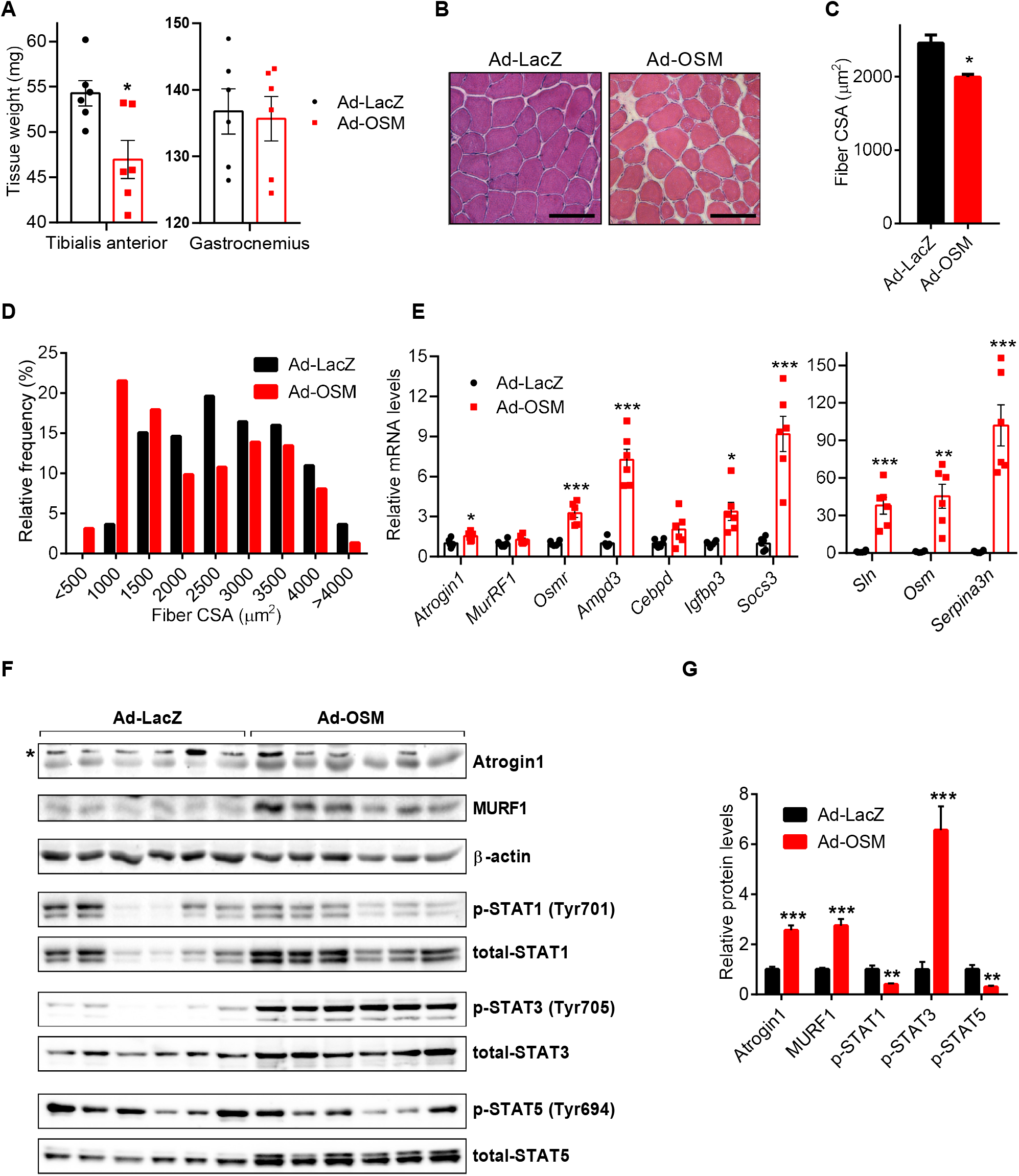
OSM overexpression causes muscle atrophy in mice. (A-G) Tibialis anterior muscles of C57BL/6 mice were transduced with LacZ or OSM expressing adenoviruses. Mice were sacrificed 7 days later (n = 6 for each group). Tissues were weighed (A) and H&E stained (B). Scale bar is 100 μm. Muscle fiber cross-sectional area (CSA) (C) and the fiber frequency distribution were determined (D) (n = 3 for each group). Changes in gene expression were determined by RT-qPCR (n = 6 for each group) (E). Protein levels were tested by western blotting. Asterisk (*) indicates non-specific band (F). Band intensities were quantified (G) (n = 6 for each group). Data are presented as mean ± SEM. Statistical analysis was conducted using two-tailed t-test. \**p* < 0.05, \*\**p* < 0.01, \*\*\**p* < 0.001, compared with the Ad-LacZ group.

### Neutralization of OSM ameliorates tumor-driven muscle wasting

We previously reported that muscle-specific OSMR knockout mice are resistant to muscle wasting. OSMR depletion in muscle tissue preserved muscle mass and strength in tumor-bearing mice (Bilgic et al., 2023). This phenotype was accompanied by reduced expression of muscle atrophy-related genes, including Atrogin1 and MURF1, suggesting that impaired OSMR activity protects from muscle loss. High plasma OSM levels were detected in tumor-bearing cachectic mice (Bilgic et al., 2023). To investigate the therapeutic potential of targeting OSM signaling to prevent cachexia-linked muscle wasting, we utilized a neutralizing anti-OSM antibody. When treated to primary myotubes along with the recombinant OSM protein, the anti-OSM antibody prevented the upregulation of OSM target genes (Figure 4A). After documenting its neutralizing effect, we administered the anti-OSM antibody to LLC-tumor bearing mice. A non-tumor-bearing group and a control tumor-bearing group received an isotype control IgG antibody. The antibody treatment was performed 10, 12, 14 and 15 days post-tumor inoculation and mice were sacrificed 1 day after the last injection (Figure 4B). Anti-OSM antibody did not affect the size of LLC tumors (Figure 4C). Remarkably, the muscle mass of the tumor-bearing mice was preserved upon anti-OSM administration (Figure 4D) and these mice exhibited improved forelimb grip strength (Figure 4E). H&E staining of gastrocnemius muscle sections also demonstrated increased muscle fiber cross-sectional area in the anti-OSM group compared to the IgG group of the tumor-bearing mice (Figure 4F and 4G). Tumor-driven increase in the frequency of muscle fibers with small cross-sectional area was suppressed upon anti-OSM administration (Figure 4H). The neutralization of OSM also reduced the phosphorylation of STAT3 and the accumulation of Atrogin1 and MURF1 proteins in muscle tissue (Figure 4I and 4J). These findings indicate that OSM plays a direct role in tumor-induced muscle wasting and the blockade of OSM activity can be used to ameliorate cachexia-associated muscle loss.

**Figure 4.**
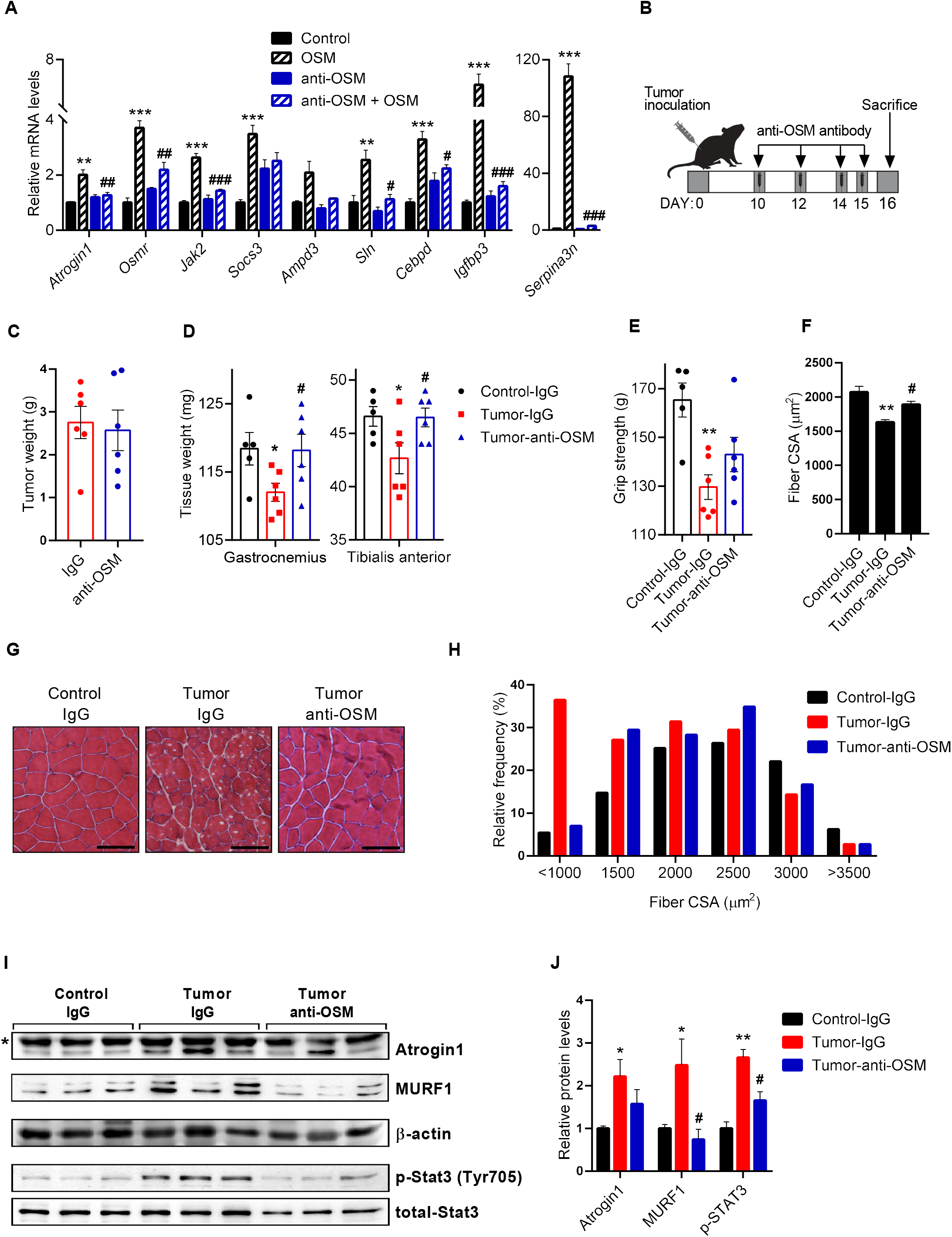
Neutralization of OSM ameliorates tumor-driven muscle wasting. (A) Mouse primary myotubes were treated with recombinant OSM (250 ng/ml) and IgG or anti-OSM antibodies (10 μg/ml) for 48 hr. Changes in gene expression were determined by RT-qPCR (n = 3 for each group). \*\**p* < 0.01, \*\*\**p* < 0.001 compares differences between IgG and IgG + OSM groups. #*p* < 0.05, ##*p* < 0.01, ###*p* < 0.001 compares differences between IgG + OSM and anti-OSM + OSM groups. (B-H) Mice inoculated with LLC cells received IgG or anti-OSM antibody injections and were sacrificed 16 days post-tumor inoculation. Tumor (C) and muscle tissues (D) were weighed. Forelimb grip strength was measured before the sacrifice (n = 5 for WT-Control group and n = 6 for other groups) (E). Gastrocnemius muscle cross-sections were H&E stained (G), cross-sectional area (CSA) (F) and the fiber frequency distribution (H) were measured (n = 3 for each group). The scale bar is 100 μm. (I-J) Gastrocnemius muscle protein levels were determined by western blotting. Asterisk (*) indicates non-specific band (G). Band intensities were quantified (J) (n = 3 for each group). \**p* < 0.05, \*\**p* < 0.01, compares differences between Control-IgG and Tumor-IgG groups. #*p* < 0.05, compares differences between Tumor-IgG and Tumor-anti-OSM groups. Data are presented as individual measurements (points) and mean ± SEM. Statistical analysis was conducted using one-way ANOVA.

### OSM target genes are upregulated in muscles of cancer cachexia and muscular dystrophy patients

We examined if the activation of OSM/OSMR signaling is linked to muscle loss in humans. We analyzed publicly available human gene expression datasets and tested transcript levels of OSM target genes. Our analysis revealed upregulated *OSMR* expression in muscle biopsies of weight-losing pancreatic ductal adenocarcinoma (PDAC) patients. Gene expression profiles of rectus abdominis muscle biopsies from 17 cachectic PDAC patients, 5 non-cachectic PDAC patients and 16 non-cancer controls were compared by *Judge et al*. using microarrays (Judge et al., 2018). Analysis of this dataset revealed upregulated *OSMR* expression in cachectic patients compared to non-cachectic and non-cancer controls (Figure 5A). We further analyzed this dataset to determine if the expression of the gene targets of OSM/OSMR signaling correlates with *OSMR* transcript levels. We compiled the top 200 genes significantly upregulated by OSM in primary myotubes and performed gene set enrichment analysis (GSEA) using this gene list. Annotated genes with known human orthologues were chosen (Table S2). GSEA analysis of the datasets revealed that OSM target genes were significantly over-represented in muscle biopsies of cachectic PDAC patients (GSE130563; normalized enrichment score (NES) = 2.17, *P-value* < 0.001) (Figure 5B).

**Figure 5.**
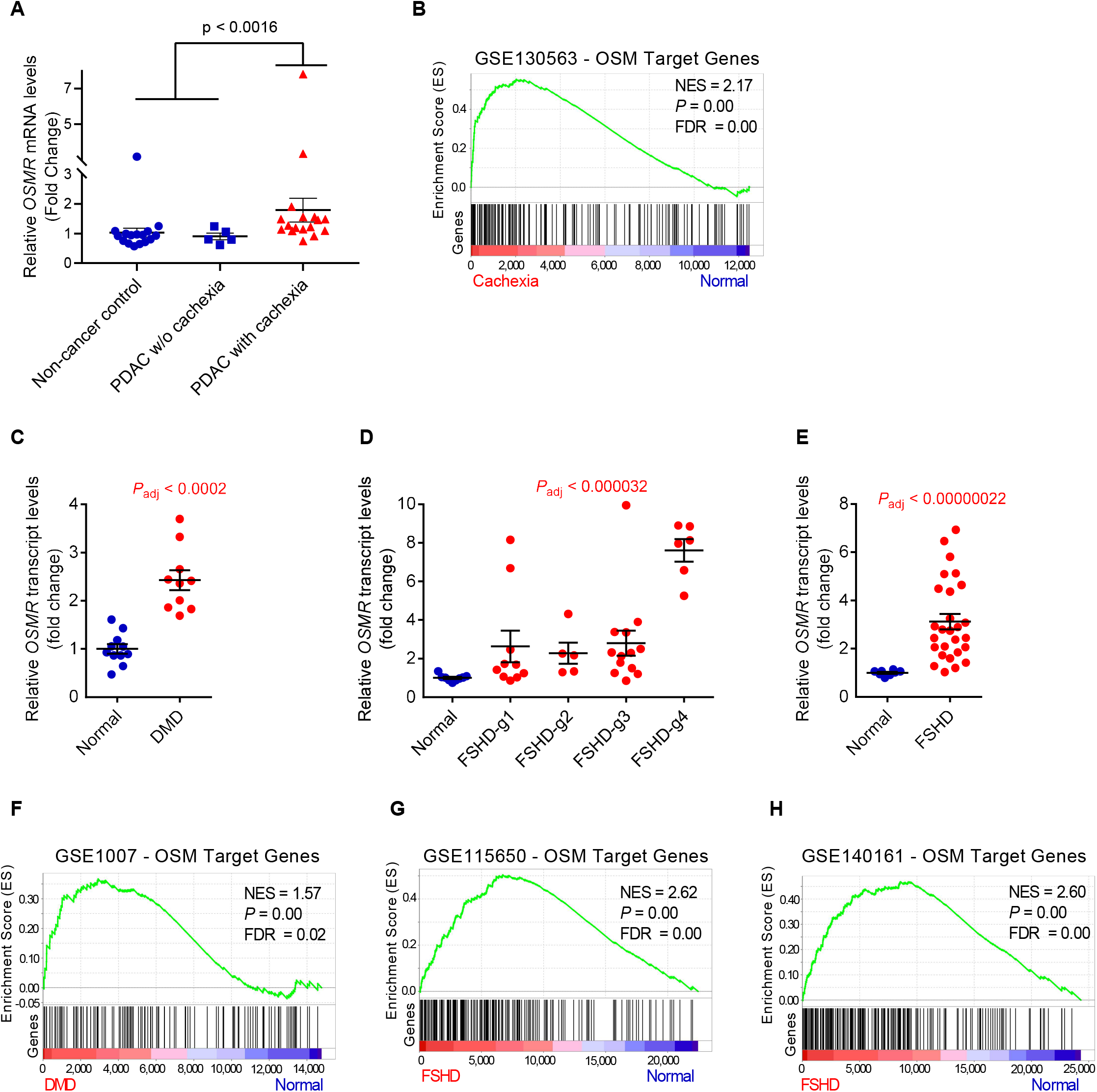
OSM target genes are upregulated in muscles of cancer cachexia and muscular dystrophy patients. (A-B) *OSMR* expression values of normal subjects and PDAC patients were analyzed by GEO2R (GSE130563) (n = 16 for non-cancer controls, n = 5 for non-cachectic PDAC patients, and n = 17 for cachectic PDAC patients) (A). Gene set enrichment analysis (GSEA) of the top 200 OSM target genes was performed by comparing cachectic PDAC patients with non-cancer controls and non-cachectic PDAC patients (GSE130563) (B). (C) *OSMR* expression values of normal subjects and DMD patients were analyzed by GEO2R (GSE1007) (n = 11 for normal subjects and n = 10 for DMD subjects). (D) *OSMR* expression values of normal subjects and FSHD patients were analyzed by DESeq2 (GSE115650) (n = 9 for normal subjects and n = 34 for FSHD subjects). (E) *OSMR* expression values of normal subjects and FSHD patients were analyzed by DESeq2 (GSE140261) (n = 8 for normal subjects and n = 27 for FSHD subjects). (F-H) Gene set enrichment analysis (GSEA) of the top 200 OSM target genes was performed. Enrichment plots for DMD (GSE1007) (F), and FSHD (GSE115650) (G), (GSE140261) (H) compared to respective normal subjects are shown. Data are presented as individual points and mean ± SEM. Statistical analysis was conducted using GEO2R or DESeq2 and values were adjusted for multiple tests.

Our gene expression analysis also revealed significantly increased *OSMR* transcript levels in muscular dystrophy diseases, including Duchenne muscular dystrophy (DMD) and Facioscapulohumeral muscular dystrophy (FSHD). Gene expression profiles of muscle biopsies from 10 DMD patients and 10 unaffected subjects were compared by Haslett *et al*. using microarrays (Haslett et al., 2003). Analysis of the datasets (GSE1007) revealed a 2.4 fold increase in *OSMR* levels in DMD patients (*P_adj_* < 0.0002) (Fig 5C). In a separate study, Dadgar *et al*. investigated 49 muscle biopsies from patients with DMD, Becker muscular dystrophy (BMD) and limb girdle muscular dystrophy (LGMD) (Dadgar et al., 2014). Analysis of this dataset (GSE109178) indicated that *OSMR* transcript was upregulated in patients with DMD (n = 17, fold change = 4.0, *P_adj_* < 0.0000009), BMD (n = 11, fold change = 3.1, *P_adj_* < 0.0022) and dysferlin-deficient LGMD2B (n = 8, fold change = 4.5, *P_adj_* < 0.009) compared to normal subjects (n = 6) (Figure S5A). The same research group also reported comparative profiling of 117 muscle biopsies from 13 different muscle disease groups. Analysis of the dataset (GSE3307) showed *OSMR* upregulation in patients with DMD (n = 10, fold change = 2.3, *P_adj_* < 0.000014), BMD (n = 5, fold change = 2.2, *P_adj_* < 0.018), acute quadriplegic myopathy (AQM) (n = 5, fold change = 2.4, *P_adj_* < 0.0019), FSHD (n = 14, fold change = 1.6, *P_adj_* < 0.008), juvenile dermatomyositis (JDM) (n = 21, fold change = 2.2, *P_adj_* < 0.0007), calpain3-deficient LGMD2A (n = 10, fold change = 2.0, *P_adj_* < 0.0007), dysferlin-deficient LGMD2B (n = 11, fold change = 2.7, *P_adj_* < 0.0007), and fokutin-related protein (FKRP)-deficient LGMD2I (n = 7, fold change = 1.7, *P_adj_* < 0.05) compared to normal subjects (n = 18) (Figure S5B and Table S3). Lastly, we analyzed the dataset generated by Wang *et al*. who collected MRI-informed muscle biopsies from FSHD patients and investigated the expression of target genes of Double homeobox 4 (DUX4) whose aberrant expression in muscle is linked to this disease (Wang et al., 2019). 9 control subjects and 36 individuals with FSHD were included in this study and patients were grouped based on expression levels of selected biomarkers. Groups 1 through 4 reflected higher DUX4 target gene expression and disease severity in ascending order with the group 4 exhibiting the highest pathology scores (Wang et al., 2019). Our analysis of the dataset (GSE115650) revealed increased *OSMR* expression in all groups with the highest levels observed in the group 4. *OSMR* was upregulated by 3.5 fold (*P_adj_* < 0.000032) in all FSHD samples combined (Figure 5D). 1-year follow up assessment of muscle biopsies was published for the same patients (GSE140261) in which *OSMR* transcript levels remained elevated (n = 27, fold change = 3.1, *P_adj_* < 0.00000022) (Figure 5E) (Wong et al., 2020).

Finally, we also tested if the OSM target genes are upregulated in muscular dystrophies. We performed GSEA analysis using the top 200 genes upregulated by OSM. Our analysis determined that OSM target genes were significantly enriched in muscle biopsies of patients with DMD (GSE1007; normalized enrichment score (NES) = 1.57, *P-value* < 0.007) (Figure 5F), and FSHD (GSE115650; NES = 2.62, *P* < 0.001 (Figure 5G) and GSE140261; NES = 2.6, *P* < 0.001) (Figure 5H). In fact, our analysis showed that the OSM target gene set is also significantly enriched in other muscular dystrophies as well, including BMD, JDM and LGMD, where high *OSMR* levels were detected (Table S3). Elevated levels of *OSMR* transcript and the enrichment of OSM target genes in muscular dystrophies implicate the activation of the OSM/OSMR pathway in these diseases.

## Discussion

In this study, we investigated the role of OSM/OSMR signaling in skeletal muscle atrophy. Our findings revealed that activation of this pathway promotes atrophy in cultured primary myotubes and in muscle tissue, and the neutralization of OSM protects from tumor-driven muscle wasting. OSM is a member of the IL-6 family of cytokines. IL-6 and LIF from the same family were previously described to stimulate muscle atrophy. Elevated IL-6 serum levels were detected in different experimental models of cancer cachexia (Baltgalvis et al., 2008; Bonetto et al., 2012) and also in cancer patients exhibiting weight loss (Scott et al., 1996). JAK/STAT3 pathway activation and muscle wasting in the presence of IL-6 were reported (Bonetto et al., 2012). The use of IL-6 receptor antibodies preserved muscle mass in IL-6 transgenic mice, mice inoculated with IL-6-overexpressing LLC cells, and in other cachexia tumor models (Ando et al., 2014; Fujita et al., 1996; Miller et al., 2017; Tsujinaka et al., 1996; White et al., 2011). Previous reports also demonstrated that LIF is secreted by cachexia-inducing C26 colon carcinoma cells and it activates the JAK/STAT3 signaling to promote atrophy in C2C12 myotubes (Seto et al., 2015). Tumor-driven LIF expression in muscle tissue was also linked to the atrophy process (Xie et al., 2021). Previously, OSM was shown to inhibit the myogenic differentiation of C2C12 cells (Xiao et al., 2011). Recently, a relevant study also reported that OSM treatment induces atrophy in C2C12 myotubes in a STAT3-dependent manner (Miki et al., 2019). Our previous work also identified OSM as an atrophy-inducing factor in primary myotubes. OSM, together with EDA-A2, additively promoted cellular atrophy and the expression of atrophy-related genes in primary myotubes (Bilgic et al., 2023). Here, we found that OSM is significantly more potent in inducing cellular atrophy compared to IL-6 and LIF. In accordance, OSM elicited more pronounced effects on the expression of atrophy-related genes in these cells.

Our results indicated that similar to IL-6 and LIF, OSM utilizes the JAK/STAT3 signaling to render its atrophic effect. STAT3 activation has been described in the atrophying muscle tissue of tumor-bearing mice. Muscle-specific STAT3 knockout mice were reported to be resistant to muscle wasting, arguing a central role for STAT3 in muscle loss (Bonetto et al., 2012; Silva et al., 2015). We found that OSM treatment of primary myotubes induces the phosphorylation of not only STAT3 but also STAT1 and STAT5. However, inhibition of STAT3 activity alone was sufficient to block OSM-driven changes in myotube diameter and atrophy-related gene expression. In fact, OSM overexpression in TA muscles of mice increased the phosphorylation of STAT3 only. Similarly, robust STAT3 activation was observed in muscles of tumor-bearing mice. Our findings implicated that the atrophy mechanism involves JAK/STAT3 signaling at the downstream of OSM/OSMR.

Here, we analyzed global gene expression profiles of primary myotubes by RNA sequencing and determined OSM target genes. In line with a previous report, we found that OSM regulated mRNA levels of *Atrogin1* but not *MuRF1* in muscle cells (Miki et al., 2019). Our analysis identified additional atrophy-related genes as transcriptional targets of the OSM pathway, such as *Serpina3n, Ampd3, Sln, Igfbp3*, and *Cepbd. Serpina3n* encodes a serine protease inhibitor, which is upregulated in injured muscle (Tjondrokoesoemo et al., 2016). *Serpina3n* expression in muscle was significantly increased in glucocorticoid-induced muscle atrophy and in C26 tumor-inoculated cachectic mice (Gueugneau et al., 2018; Shum et al., 2015). Serpina3n was proposed as a circulating biomarker of muscle atrophy (Gueugneau et al., 2018). In fact, we found that *Serpina3n* is one of the top gene targets of the OSM pathway in myotubes. Ampd3 controls adenine nucleotide content to ATP ratio and plays an important role in balancing muscle metabolism (Miller et al., 2019). *Ampd3* levels were upregulated in muscles undergoing atrophy (Brocca et al., 2017; Ibebunjo et al., 2013; Lecker et al., 2004). Its overexpression in C2C12 cells or TA muscles of mice was shown to mimic metabolic changes that occur during muscle atrophy (Miller et al., 2021). *Sln* (sarcolipin) encodes a micropeptide which regulates the activity of sarcoplasmic reticulum Ca^2+^-ATPase pumps. *Sln* expression is upregulated in several myopathies including DMD (Schneider et al., 2013). Knock-down of *Sln* ameliorated muscle physiology and survival in a mouse model of DMD (Voit et al., 2017). *Igfbp3* encodes an IGF-binding protein, which was shown to suppress IGF signaling and promote atrophy in C2C12 myotubes (Huang et al., 2016). Increased *Igfbp3* levels were detected in muscles of tumor-bearing mice (Cole et al., 2021). *Cebpd* (CAAT/enhancer-binding protein δ) expression was induced in atrophying muscles of glucocorticoid-treated and tumor-bearing mice (Fontes-Oliveira et al., 2014; Yang et al., 2005). STAT3-induced *Cebpd* activity was shown to promote Myostatin and Atrogin1 expression in muscle and *Cebpd* deletion in mice protected from tumor-induced muscle wasting (Silva et al., 2015; Zhang et al., 2013). Our results indicated that transcription of these atrophy-related genes is regulated by OSM in a JAK/STAT3-dependent manner. Further investigation is needed to assess the potential contribution of these target genes to OSM-induced muscle atrophy.

Evidence supporting the role of OSM in cancer cachexia comes from tumor inoculation studies utilizing muscle-specific OSMR-knockout mice (Bilgic et al., 2023). Upon tumor growth, these mice exhibited reduced muscle wasting and improved muscle strength compared to their wild-type controls. Expression of muscle atrophy-related genes, including *Atrogin1, MuRF1* and *Eda2r*, was reduced in muscles of tumor-bearing OSMR-deficient mice (Bilgic et al., 2023). The result presented here corroborates these findings and highlights the therapeutic potential of blocking the OSM/OSMR signaling. The neutralization of tumor-induced OSM using a specific antibody phenocopied the OSMR-deficiency in mice. Our results indicated that OSM blockade in tumor-bearing mice attenuates muscle atrophy and preserves muscle strength, accompanied by reduced expression of atrophy genes in muscle tissue. Previously, IL-6 was reported as the main inflammatory mediator driving muscle wasting during cancer cachexia. However, anti-IL-6 therapies did not satisfactorily prevent muscle wasting in cancer patients (Bayliss et al., 2011; Marceca et al., 2020). A previous tumor inoculation study utilizing IL-6 receptor knockout mice showed preservation of fat mass without an effect on muscle wasting (Petruzzelli et al., 2014). Our results argue that OSMR protein is necessary for tumor-driven muscle loss and the inhibition of OSM/OSMR signaling may be useful in preventing cancer cachexia. Future studies should utilize neutralizing antibodies against human OSM/OSMR to establish a clinical benefit. We detected elevated transcript levels of *OSMR* and other OSM targets in muscle biopsies of PDAC patients exhibiting cachexia. The patient groups that would maximally benefit from an OSM/OSMR therapy should be determined by testing for elevated OSM levels in tumor and plasma samples.

Lastly, our analysis of publicly available gene expression datasets showed that *OSMR* transcript levels were significantly upregulated and OSM target genes were significantly enriched in muscle biopsies of patients with muscular dystrophies, such as DMD and FSHD. These results hinted at the activation of OSM/OSMR signaling in these muscle samples. In fact, inflammatory response has been indicated to play a central role in the progression of muscular dystrophies (Raimondo and Mooney, 2021; Rosenberg et al., 2015). Therefore, it is tempting to ask whether OSM functions as an inflammatory mediator involved in the development of muscle weakness and damage in muscular dystrophies. Further studies utilizing animal models and testing protein-level changes in patient samples are required to establish a role for the OSM/OSMR pathway in muscle loss associated with these diseases.

## Acknowledgements

We gratefully acknowledge the use of the Koc University Research Center for Translational Medicine (KUTTAM) animal facility infrastructure. We would like to thank Prof. Gerhard Müller-Newen (RWTH Aachen University) for providing the mouse STAT3-Y705F plasmid. Also, we greatly appreciate help from Beril Esin for her support in the GEO database search. This work was supported by the EMBO installation grant (#4162) and The Scientific and Technological Research Council of Türkiye (TUBITAK) grants 118Z791 and 118C014 to S. Kir

## Author Contributions

S.K. conceived and designed the experiments. A.D., S.A., B.T., A.E.K., S.N.B. and S.K. performed the experiments. A.D., S.A., B.T., A.E.K. and S.K. analyzed the data. A.D. and S.K. wrote the manuscript.

## Materials and methods

### Reagents

Recombinant proteins OSM (495-MO), IL-6 (406-ML) and LIF (8878-LF) were purchased from R&D Systems. JAK1/2 inhibitor Ruxolitinib was obtained from (Abcam; ab141356). Mouse OSM expression plasmid (MR226014) was purchased from Origene. Mouse STAT3-Y705F plasmid was kindly provided by Prof. Gerhard Müller-Newen (RWTH Aachen University). Mouse OSM antibody (AF-495-NA) and control goat IgG antibody (AB-108-C) were purchased from R&D Systems.

### Mice

All experimental procedures were conducted in the Koc University Animal Research Facility in accordance with institutional policies and animal care ethics guidelines. 8-12-week-old male mice were housed in 12-hour light/dark cycles (7am-7pm) and provided ad libitum access to standard rodent chow diet and water. All mice were maintained on a pure C57BL/6 background. All animal protocols were approved by the Institutional Animal Care and Use Committee of Koc University.

### Tumor inoculation and antibody administration

LLC cells were cultured in DMEM medium (Sigma 5796) supplemented with 10% Fetal Bovine Serum (FBS) and 1 % penicillin/streptomycin (Invitrogen). Mice were randomly assigned into groups while satisfying the criteria that the average body weight in each group is similar. All mice used in tumor inoculation experiments were from C57BL/6 background. LLC cells (5 × 10^6^ per mouse) cells were injected subcutaneously over the flank. Non-tumor-bearing control mice received the vehicle (PBS) only. Mice were housed individually in all tumor inoculation experiments. In antibody administration experiments, mice were injected with control IgG (AB-108-C) or anti-OSM (AF-495-NA) antibodies (1 mg/kg) for 4 times; 10, 12, 14 and 15 days post-tumor inoculation. Mice were sacrificed 16 days after tumor inoculation. Gastrocnemius and tibialis anterior muscles and tumors were dissected and weighed using an analytical balance.

### Grip strength

Forelimb grip forces were measured on the same day as sacrifice using grip strength meter (Ugo Basile grip strength meter). Each mouse was held from its tail and allowed to grab a bar attached to a force transducer while being pulled horizontally away from the bar. The peak force applied before releasing the bar was registered from at least 3 repetitions and averaged to determine the grip strength of each mouse.

### Tissue histology

Muscle tissues were frozen in liquid nitrogen-cooled isopentane for 15-20 seconds. Frozen tissues were embedded in cryomolds in Tissue-Tek OCT freezing medium (Sakura). For hematoxylin & eosin staining, 7 μm thick sections were cut using a cryostat (Leica) and collected on Superfrost Plus slides (Thermo). Sections were fixed with 4% paraformaldehyde and treated with hematoxylin (Merck 105174), 0.1% HCl, eosin (Merck 109844), 70-100% ethanol gradient and xylene (Isolab), respectively. The muscle fiber cross-sectional area was assessed using Image J software.

### Adenovirus production and injection

Adenovirus vectors were generated using the Virapower Adenoviral expression system (Invitrogen). Open reading frames of *Osm* and *Stat3-Y705F* following a CACC sequence were cloned into a pENTR-D-TOPO plasmid and then recombined into the pAd-CMV-DEST adenoviral plasmid using LR clonase II. LacZ adenoviral plasmid was provided in the kit. After digestion with PacI (Thermo), adenoviral plasmids were transfected into 293A cells using Lipofectamine 2000 (Invitrogen). 293A cells were cultured in DMEM (Sigma 5796), 10% FBS (Invitrogen) and penicillin/streptomycin (Invitrogen). Adenoviral particles were collected from cell culture supernatant following the manufacturer’s instructions. Adeno-X Maxi Purification Kit and Adeno-X Rapid Titer Kit from Clontech were used for purification and titration, respectively. Mice were injected unilaterally with 5×10^8^ infectious units of Adeno-OSM into the tibialis anterior muscle while the contralateral muscle received the same dose of control Adeno-LacZ. Mice were sacrificed 7 days later.

### Myoblast culture

Mouse primary myoblasts were isolated from limb muscles of pups (2-3 days old) and cultured in Ham’s F-10 nutrient mixture (Invitrogen) with 20% FBS (Invitrogen) supplemented with 2.5 ng/ml basic fibroblast growth factor (bFGF) (Sigma) and penicillin/streptomycin (Invitrogen). Myoblasts were then transferred to DMEM (Sigma 5796) supplemented with 5% horse serum and penicillin/streptomycin (Invitrogen) for differentiation. Myotube cells were harvested within 72 hr of differentiation. Adeno-GFP was added to the cells at the start of differentiation for fluorescent myotube imaging performed using a live cell imager (Zeiss Axiolab live-cell imager). The diameters of individual myotubes were measured using Image J software. Treatments with other adenoviruses were performed 24 hr after differentiation.

### Western blotting

Total protein from cells was isolated using lysis buffer containing 50 mM Tris (pH 7.5), 150 mM NaCl, 1% Triton X-100, 5 mM EDTA, 1 mM PMSF, supplemented with protease inhibitor tablets (Roche) and phosphatase inhibitors; 20 mM NaF, 10 mM β-glycerol phosphate, 10 mM Na_4_P2O_7_, 2 mM Na_3_VO_4_. A similar lysis buffer was used for tissue samples where 1% NP40 was used as the detergent and 10% glycerol was added. Tissues were homogenized using a Kinematica (PT1200E) homogenizer. The homogenates were centrifuged at 13,000 rpm for 10 min and the supernatants were used as lysates. Protein concentration was determined by Bio-Rad Protein assay. For each run, 30 μg of protein lysate was resolved in SDS-polyacrylamide gel and then blotted in a nitrocellulose membrane (Thermo). After blocking with 5% non-fat milk in Tris-buffered saline containing 0.05% Tween 20 (TBS-T), the membrane was blotted overnight with primary antibodies in TBS-T containing 5% BSA (Cell signaling antibodies; Stat1 (9172), phospho-Stat1-Tyr701-(7649), Stat3 (9139), phospho-Stat3-Tyr705 (9145), Stat5 (94205T), phospho-Stat5-Tyr694 (4322T), Jak2 (3230), phospho-Jak2-Tyr1008 (8082), β-actin (3700), p65-RelA (8242), phospho-p65-RelA-Ser536 (3033), p105/p50-NFκB (12540), phospho-p105-NFκB-Ser932 (4806), p100/p52-NFκB2 (4882), IκB (4814), phospho-IκB-Ser32 (2859), ECM Bioscience antibodies; Atrogin1 (AP2041) and MURF1 (MP3401), and DSHB; MyHC antibody (MF20)). For secondary antibody incubation, TBS-T containing 5% milk was used (Cell signaling, anti-rabbit (7074) and anti-mouse (7076)). WesternBright blotting substrates from Advansta were used to visualize the results on a Chemidoc imaging system (Bio-rad). For blots visualized with a Licor Odyssey CLx imaging system, IRDye 680RD anti-mouse (926-68070) and IRDye 800CW anti-rabbit (926-32211) secondary antibodies were used. Blot images were quantified using Image J Software. Phosphorylated proteins were normalized to their respective total protein signals. Atrogin1 and MURF1 levels were normalized to a non-specific band or β-actin signal.

### Immunofluorescence

Primary myotubes were fixed with pre-chilled methanol for 10min at −20°C, incubated in a blocking solution (3% bovine serum albumin + 0.1% Triton X-100 + 10% Horse serum) for 1hr at room temperature and then incubated in the blocking solution containing myosin heavy chain (MyHC) antibody (1:1000) (DSHB, MF20) for 1hr. Myotubes were washed with PBS and incubated with anti-mouse IgG H&L Alexa Fluor 594 secondary antibody (1:2000) (Abcam, ab150116) and DAPI (1:3000) (Cayman, 14285) in the blocking solution for 1hr at room temperature and then mounted using homemade mounting medium. Cells were visualized using fluorescence microscopy (Zeiss). MyHC signal was normalized to the number of myotube nuclei. Fields with similar density of myotubes were chosen.

### RT-qPCR

Total RNA from cultured cells or tissue samples was extracted using Qiazol reagent (Qiagen) and purified with RNA spin columns (Ecotech). Tissues were homogenized using TissueLyzer LT (Qiagen). Complementary DNA synthesis was carried out with High-Capacity cDNA Reverse Transcription kit (Thermo). The resultant cDNA was analyzed by RT–qPCR using a CFX Connect instrument (Bio-Rad). In each reaction, 25 ng of cDNA and 150 nmol of each primer were mixed with iTaq Universal SYBR Green Supermix (Bio-Rad). Relative mRNA levels were calculated by the ΔΔCt method and normalized to cyclophilin mRNA. The following primers were used: *Cyclo* F: 5’-GGAGATGGCACAGGAGGAA-3’, R: 5’-GCCCGTAGTGCTTCAGCTT-3. *Actb* F: 5’-TTCTTGGGTATGGAATCCTGTGG-3’, R: 5’-TTTACGGATGTCAACGTCACAC-3’. *Atrogin1* F: 5’-TCAGAGAGGCAGATTCGCAA-3’, R: 5’-GGGTGACCCCATACTGCTCT-3’. *MuRF1* F: 5’-TCCTGATGGAAACGCTATGGAG-3’, R: 5’-ATTCGCAGCCTGGAAGATGT-3’. *Osmr* F: 5’-TCACAACTCCAGATGCACGC-3’, R: 5’-ACTTCTCCTTCACCCACTGAC-3’. *Jak2* F: 5’-CCACCCGTGGAATTTATGCG −3’, R: 5’-TGGCAATCTTCCGTTGCTCT −3’. *Socs3* F: 5’-TAGACTTCACGGCTGCCAAC −3’, R: 5’-CGGGGAGCTAGTCCCGAA-3’. *Ampd3* F: 5’-CTCCTCTCAGCAACAACAGCC-3’, R: 5’-CTCCATGAGCGCTTCCTTTGTG-3’. *Sln* F: 5’-GGTGGAGAGACTGAGGTCCTT-3’, R: 5’-CCAAGGCTTGTCTTCACTTCCTGA-3’. *Serpina3n* F: 5’-AGAGTCTCTCAGGTGGTCC-3’, R: 5’-ACAGTTTCGCAGACATTGGGA-3’. *Stat3* F: 5’-AGGAGTCTAACAACGGCAGC-3’, R: 5’-GTCACGATCAAGGAGGCATCA-3’. *Igfbp3* F: 5’-ATAAGAAGAAGCAGTGCCGCC-3’, R: 5’-GTCGTCTTTCCCCTTGGTGT-3’. *Cebpd* F: 5’-AGAACCCGCGGCCTTCTAC-3’, R: 5’-ATGTAGGCGCTGAAGTCGAT-3’. *Osm* F: 5’-GAACACTGCTCAGTTTGACCC-3’, R: 5’-CGTGAGGTTCGCCTGATTCT-3’.

### RNA sequencing

Differentiated mouse primary myotubes were treated with OSM, LIF or IL6 for 48 hr (250 ng/ml each). Total RNA was isolated and DNase-treated using Qiazol reagent (Qiagen) and Direct-zol RNA MiniPrep kit (Zymo Research). Samples were prepared using Illumina TruSeq Stranded mRNA Library Prep Kit. Pair-end sequencing was performed using Illumina NovaSeq technology producing a total of 60 million (2 x 100bp) paired-end reads per sample. The quality check of the raw and trimmed data was performed with FastQC (v.0.11.9) and MultiQC (v1.12). Adaptor sequences and poor-quality sequences were removed with trimGalore (v0.6.5). Mouse genome index (GRCm39) was created with STAR (v2.7.3a) using genomeGenerate run mode. Trimmed sequences were mapped onto this index using STAR alignReads run mode. Quality check of alignments was performed with qualimap (v.2.2.1). Alignments were quantified using HTSeq-count (v0.11.1) Python package. Differential gene expression analysis was done with R Studio (v2021.09.2+382) and DESeq2 (v1.34.0) R package. Plots were visualized with ggplot2 (v3.3.5) and EnhancedVolcano (v.1.12.0) R packages. Gene set enrichment analysis results and plots were generated using GSEA (v4.1.0) software.

### Human gene expression analysis

Gene expression profiles of skeletal muscle samples from PDAC patients, muscular dystrophy patients and normal subjects were accessed from Gene Expression Omnibus (GEO) database as normalized datasets. Microarray datasets (GSE130563, GSE1007, GSE109178 and GSE3307) were analyzed by GEO2R using default settings. RNA sequencing datasets (GSE115650 and GSE140261) were also accessed from the GEO database. Gene count normalization and fold change calculations were performed using the DEseq2 R package as described above. *P* values were adjusted for multiple testing. Gene set enrichment analysis was performed using GSEA (v4.1.0) software. Microarray data provided in multiple files (GSE1007 and GSE3307) were combined sample-wise to create a single dataset for GSEA analysis.

### Statistical Analysis

Values are expressed as mean ± SEM. Error bars (SEM) shown in all results were derived from biological replicates. Comparisons between two groups were evaluated using a two-tailed, unpaired *t*-test. Comparisons of more than two groups were performed using one-way ANOVA and corrected for multiple comparisons using Tukey’s post-hoc test.

### Lead Contact

Further information and requests for resources and reagents should be directed to and will be fulfilled by the Lead Contact, Serkan Kir.

### Materials Availability

All unique/stable reagents generated in this study are available from the Lead Contact with a completed Materials Transfer Agreement.

### Data Availability

RNA sequencing data generated in this study are available in the GEO database with accession number GSE222208. Human gene expression datasets analyzed here are also available in the GEO database; GSE130563 (Judge et al., 2018), GSE1007 (Haslett et al., 2003), GSE109178 (Dadgar et al., 2014), GSE115650 (Wang et al., 2019), GSE140261 (Wong et al., 2020), and GSE3307 (Dadgar et al., 2014).

## Supplemental Information

**Figure S1.**
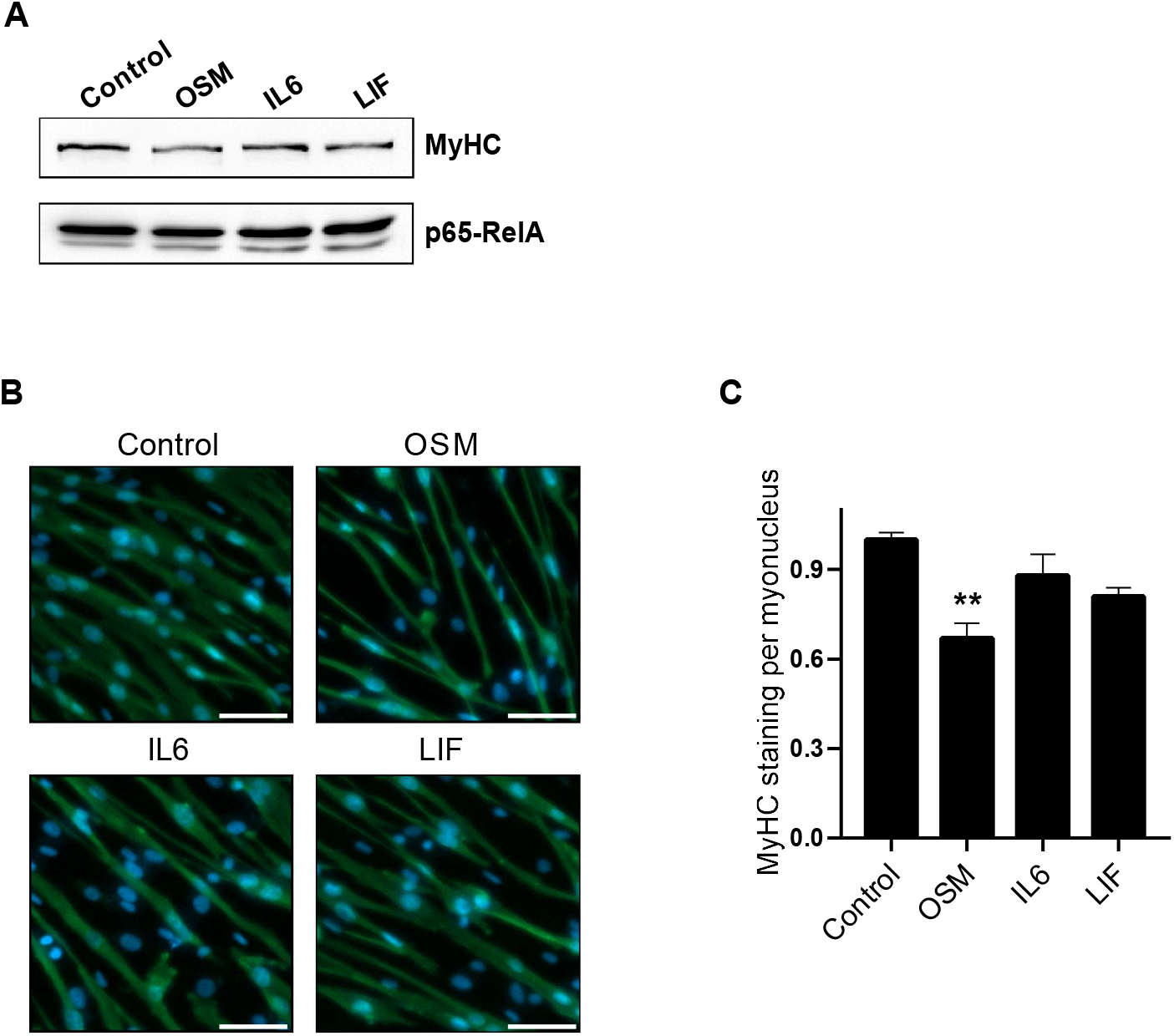
OSM promotes atrophy in cultured primary myotubes. (A-C) Primary myotubes were treated with OSM, IL6 and LIF (each 250 ng/ml) for 48 hr. Protein levels were determined by western blotting (A). Cells were also immunofluorescently labelled for MyHC and their nuclei were counterstained with DAPI. The scale bar is 50 μm (B). MyHC signal was normalized to the number of myotube nuclei (n = 4) (C).

**Figure S2.**
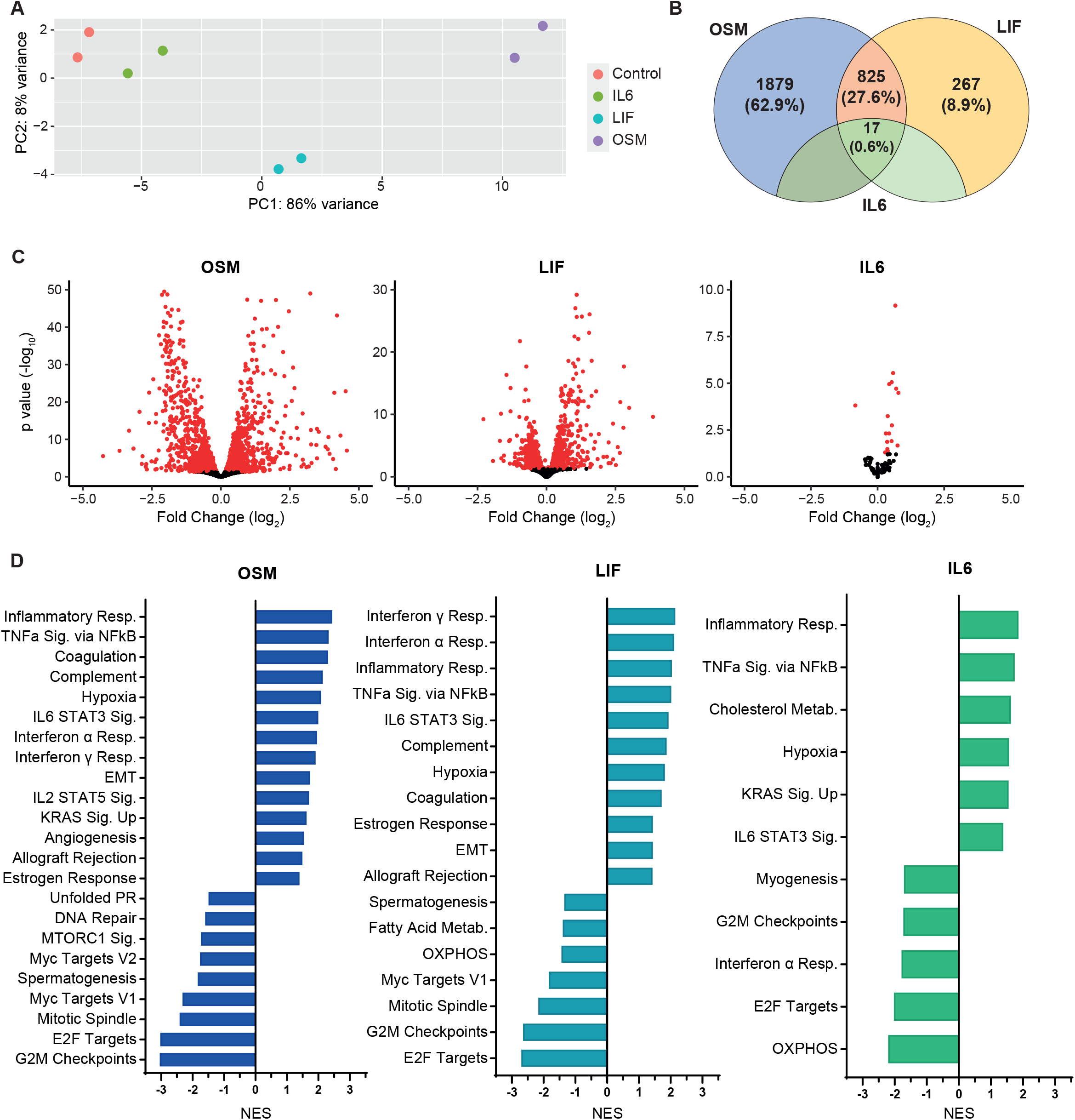
RNA sequencing analysis of primary myotubes treated with the IL6 family cytokines. (A) RNA sequencing dataset was analyzed using DESeq2. The principal component analysis is shown. The separation between gene expression profiles is depicted. (B) The number of significantly differentially expressed genes for OSM, LIF, IL6 and their intersections is shown. (C) Volcano plots of RNA sequencing analysis are shown. Significantly differentially expressed genes are represented by red dots. (D) Gene ontology analysis of the RNA sequencing data was performed. Significantly up-and down-regulated pathways are shown. NES, normalized enrichment score. RNA sequencing count data was normalized and fold change calculations were performed using the DEseq2 R package. *P* values were adjusted for multiple tests. Differentially expressed genes with P_adj_ < 0.05 were chosen.

**Figure S3.**
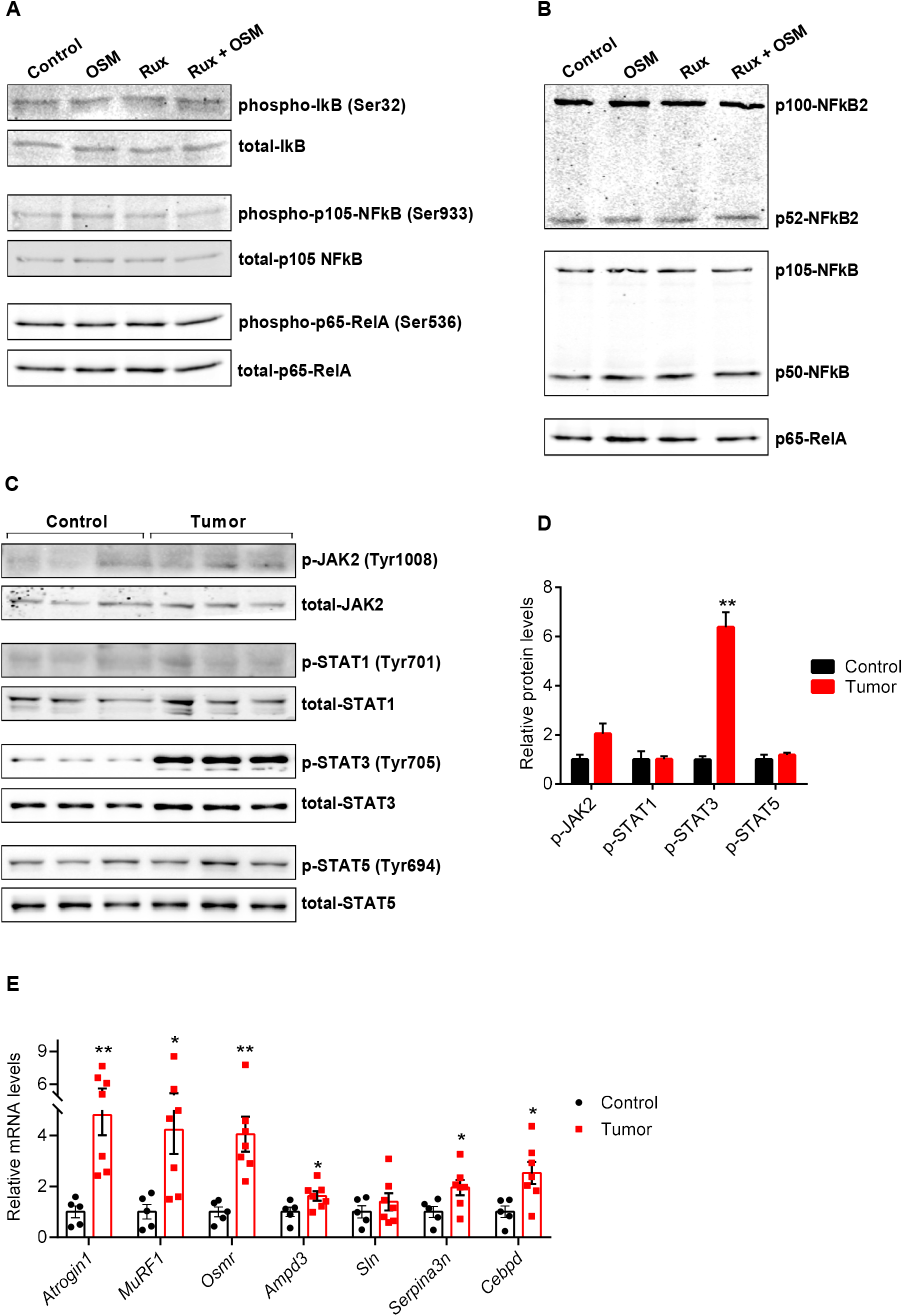
OSM protein does not activate the NFκB signaling in primary myotubes and LLC tumors induce JAK/STAT3 signaling and OSM target gene expression in gastrocnemius muscles of mice. (A) Mouse primary myotubes were treated with Ruxolitinib (2 μM) for 30 min and then recombinant OSM (250 ng/ml) was added for 10 min. Protein levels were determined by western blotting. (B) Mouse primary myotubes were treated with Ruxolitinib (2 μM) and recombinant OSM (250 ng/ml) for 24 hr. Protein levels were determined by western blotting. (C-E) C57BL/6 mice inoculated with LLC cells were sacrificed 16 days later. Protein levels in gastrocnemius muscle were determined by western blotting (C) and quantified (D) (n = 3 for each group). mRNA levels in gastrocnemius muscle were tested by RT-qPCR (n = 5 for the control group and n = 7 for the tumor group) (E). Data are presented as mean ± SEM. Statistical analysis was conducted using a two-tailed *t*-test. \**p* < 0.05, \*\**p* < 0.01, compared with the control group.

**Figure S4.**
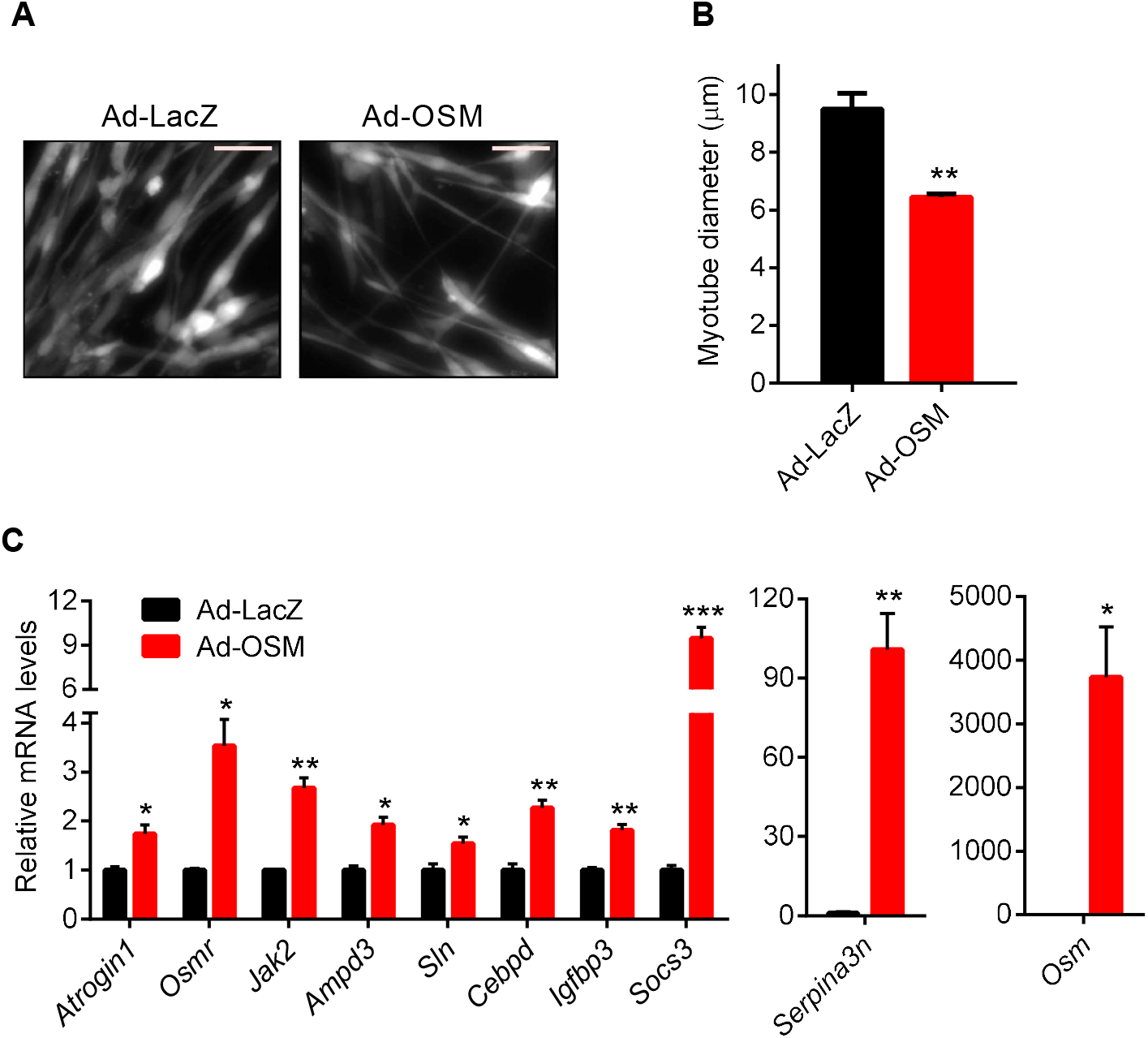
OSM overexpression promotes cellular atrophy in primary myotubes. (A and B) Mouse primary myotubes were transduced with LacZ or OSM expressing adenoviruses and examined 48 hr later. Cells were also transduced with a GFP adenovirus and visualized under the fluorescence microscope. The scale bar is 50 μm (A). Average myotube diameter was measured (n = 4 for each group) (B). (C) Mouse primary myotubes were transduced with LacZ or OSM expressing adenoviruses and harvested 48 hr later. mRNA levels were determined by RT-qPCR (n = 3 for each group). Data are presented as mean ± SEM. Statistical analysis was conducted using two-tailed *t*-test. \**p* < 0.05, \*\**p* < 0.01, \*\*\**p* < 0.001, compared with the Ad-LacZ group.

**Figure S5.**
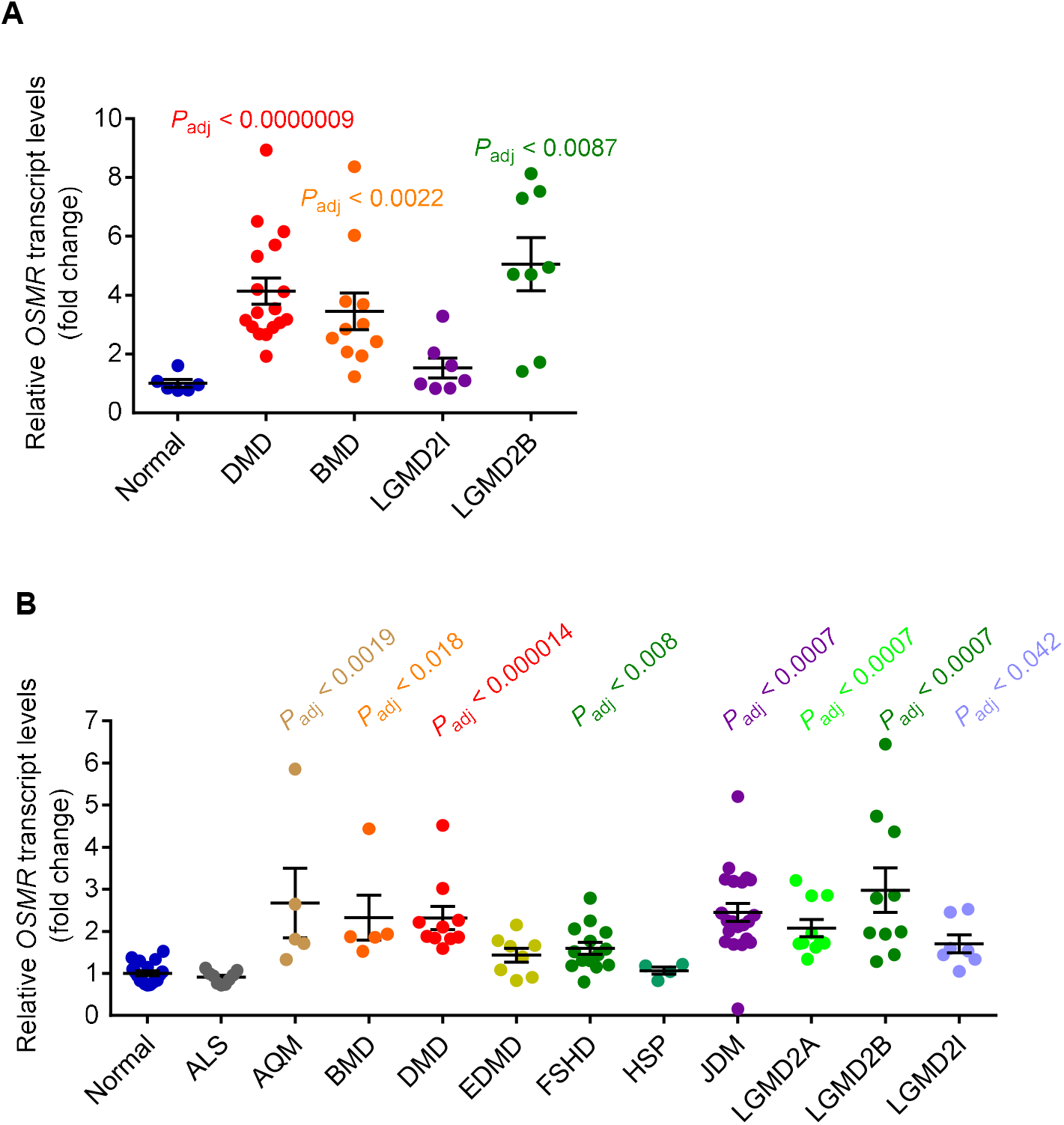
*OSMR* is upregulated in muscle biopsies of muscle dystrophy patients. (A) *OSMR* expression values of normal subjects and DMD patients were analyzed by GEO2R (GSE109178) (n = 6 for normal subjects, n = 17 for DMD subjects, n = 11 for Becker muscular dystrophy (BMD) subjects, n = 7 for fukutin-related protein (FKRP)-deficient limb girdle muscular dystrophy type 2I (LGMD2I) subjects and n = 8 for dysferlin-deficient limb-girdle muscular dystrophy type 2B (LGMD2B) subjects). (B) *OSMR* expression values of normal subjects and muscular dystrophy patients were analyzed by GEO2R (GSE3307) (n = 18 for normal subjects, n = 9 for amyotrophic lateral sclerosis (ALS) patients, n = 5 for acute quadriplegic myopathy (AQM), n = 5 for Becker muscular dystrophy (BMD) patients, n = 10 for DMD patients, n = 8 for Emery Dreifuss muscular dystrophy (EDMD) patients, n = 14 for FSHD patients, n = 4 for hereditary spastic paraplegia (HSP) patients, n = 21 for juvenile dermatomyositis (JDM) patients, n =10 for calpain3-deficient limb girdle muscular dystrophy type 2A (LGMD2A) patients, n = 10 for dysferlin-deficient limb girdle muscular dystrophy type 2B (LGMD2B) patients, and n = 7 for fukutin-related protein (FKRP)-deficient limb girdle muscular dystrophy type 2I (LGMD2I) patients). Data are presented as individual points and mean ± SEM. Statistical analysis was conducted using GEO2R and values were adjusted for multiple tests.

**Table S1. The list of differentially expressed genes.**

Differentially expressed genes with *P*_adj_ < 0.05 identified in RNA sequencing of mouse primary myotubes treated with recombinant OSM, LIF, or IL6 proteins (250 ng/ml each; 48hr) are shown.

**Table S2. The list of human orthologues of the top 200 OSM target genes.**

The top 200 significantly upregulated genes in primary myotubes in response to OSM treatment are listed. Annotated genes with known human orthologues were chosen.

**Table S3. Analysis of *OSMR* transcript levels and OSM target gene set enrichment in muscular dystrophy datasets.**

Fold change (FC) and adjusted *P* values (*P*_adj_) of *OSMR* transcript levels compared to normal subjects are listed. Normalized enrichment scores (NES), *P* values (representing false discovery rate), and the number of genes present in the dataset (set size) are shown for GSEA analysis testing enrichment of OSM target genes.

